# The effective population size modulates the strength of GC biased gene conversion in two passerines

**DOI:** 10.1101/2021.04.20.440602

**Authors:** Henry J Barton, Kai Zeng

## Abstract

Understanding the determinants of genomic base composition is fundamental to understanding genome evolution. GC biased gene conversion (gBGC) is a key driving force behind genomic GC content, through the preferential incorporation of GC alleles over AT alleles during recombination, driving them towards fixation. The majority of work on gBGC has focussed on its role in coding regions, largely to address how it confounds estimates of selection. Non-coding regions have received less attention, particularly in regard to the interaction of gBGC and the effective population size (*N_e_*) within and between species. To address this, we investigate how the strength of gBGC (*B* = 4*N_e_b*, where *b* is the conversion bias) varies within the non-coding genome of two wild passerines. We use a dataset of published high coverage genomes (10 great tits and 10 zebra finches) to estimate *B*, nucleotide diversity, changes in *N_e_*, and crossover rates from linkage maps, in 1Mb homologous windows in each species. We demonstrate remarkable conservation of both *B* and crossover rate between species. We show that the mean strength of gBGC in the zebra finch is more than double that in the great tit, consistent with its twofold greater effective population size. *B* also correlates with both crossover rate and nucleotide diversity in each species. Finally, we estimate equilibrium GC content from both divergence and polymorphism data, which indicates that *B* has been increasing in both species, and provide support for population expansion explaining a large proportion of this increase in the zebra finch.

**Significance statement:** Understanding the forces that change the nucleotide base composition of genomes is central to understanding their evolution. One such force is GC biased gene conversion, a process that during recombination converts some heterozygous base positions to homozygous. This process is more likely to convert adenine and thymine bases to guanine and cytosine bases than the other way around, hence is GC biased. This increases the frequency of GC alleles in a way similar to positive selection. This process has largely been studied within protein coding regions, and not often compared between species. We measure its strength in the non-coding areas of the genomes of two bird species, showing it to be stronger in the species with the larger population size.

## Introduction

A large proportion of many organisms’ genomes are non-coding; 99% in humans, 80% in *Drosophila melanogaster*, 73% in *Caenorhabditis elegans* and 71% in *Arabidopsis thaliana* (Halligan and Keightley, 2006; Rajic *et al.*, 2005). The non-coding genome offers the opportunity to study evolutionary process away from the interference of the direct effects of natural selection. One such process is the evolution of genomic base composition. The evolution of base content and its variation within genomes has been the focus of intrigue for many years, such as the question of mammalian isochore evolution (Eyre-Walker and Hurst, 2001). Genomic GC content is predominately determined by the balance between the strong (G and C bases) to weak (A and T bases) substitution rate (S→W), in part underpinned by CpG hypermutabiliy (Hodgkinson and Eyre-Walker, 2011; Hwang and Green, 2004; Ségurel *et al.*, 2014), and the weak to strong substitution rate (W→S), which is influenced by GC biased gene conversion (gBGC), which favours strong over weak bases, and is a major determinant of GC content evolution in a broad range of organisms (Bolívar *et al.*, 2016, 2018, 2019; Corcoran *et al.*, 2017; Glémin *et al.*, 2015; Gossmann *et al.*, 2018; Jackson *et al.*, 2017; Muyle *et al.*, 2011; Ratnakumar *et al.*, 2010; Wallberg *et al.*, 2015). Although, recent experimental based measures of gene conversion in *Saccharomyces cerevisiae*, *Neurospora crassa*, *Chlamydomonas reinhardtii* and *Arabidopsis thaliana*, did not reveal a conversion bias (Liu *et al.*, 2018).

gBGC is the preferential incorporation of GC alleles over AT alleles during the resolution of heteroduplex DNA resulting from the repair of double stranded breaks during recombination (Chen *et al.*, 2007; Duret and Galtier, 2009). This elevates the number of gametes containing GC alleles, as observed in humans (Williams *et al.*, 2015) and birds (Smeds *et al.*, 2016). As such, gBGC acts to increase the frequency of G and C alleles over A and T alleles, in a manner that mirrors positive selection (Duret and Galtier, 2009; Galtier and Duret, 2007; Gutz and Leslie, 1976; Nagylaki, 1983). As a result, gBGC is an inconvenient complication when looking for signatures of selection in genomes. For example, over 20% of identified positively selected genes in the human lineage are possibly just the focus of elevated gBGC (Ratnakumar *et al.*, 2010). Furthermore, a growing body of literature has demonstrated that gBGC confounds our ability to estimate parameters such as the rate of adaptation (*ω* = *dN/dS*) (Bolívar *et al.*, 2018, 2019; Corcoran *et al.*, 2017; Gossmann *et al.*, 2018; Ratnakumar *et al.*, 2010; Rousselle *et al.*, 2019) and the proportion of substitutions fixed by positive selection (*α*) (Bolívar *et al.*, 2018; Corcoran *et al.*, 2017; Rousselle *et al.*, 2019). Equally, studying gBGC in coding regions is inconvenienced by the action of natural selection also acting on those regions, forcing studies to use putatively neutral sites like third codon positions (Rousselle *et al.*, 2019; Weber *et al.*, 2014) and 4-fold degenerate sites (Bolívar *et al.*, 2016; Corcoran *et al.*, 2017; Gossmann *et al.*, 2018) reducing the amount of data available as well as potentially being confounded by codon usage bias (Chamary and Hurst, 2005; Galtier *et al.*, 2018; Hayes *et al.*, 2020; Jackson *et al.*, 2017; Kunstner *et al.*, 2011).

As gBGC is a recombination mediated process, it should co-vary in strength with crossover rate, at different genomic scales and between species. This is seen in a large body of literature, demonstrating correlations between recombination rate and GC content (Bolívar *et al.*, 2016; Glémin *et al.*, 2015; Rousselle *et al.*, 2019; Wallberg *et al.*, 2015; Weber *et al.*, 2014), recombination rate and equilibrium GC content (GC*) (Duret and Arndt, 2008; Muyle *et al.*, 2011; Singhal *et al.*, 2015), and recombination rate and the population scaled strength of gBGC, *B* = 4*N_e_b*, where *N_e_* is the effective population size and *b* is the raw strength of conversion bias (Glémin *et al.*, 2015; Wallberg *et al.*, 2015). However, notably, in *Dropshophila* gene conversion rate does not positively correlate with crossover rate (Comeron *et al.*, 2012). With recombination varying greatly between organisms (Stapley *et al.*, 2017), gBGC can also be expected to have similar variation in strength and impact. For example, in mammals the recombination landscape is largely determined by the location of recombination hotspots, determined by the PRDM9 gene (Baudat *et al.*, 2010; Parvanov *et al.*, 2010). This results in areas of greatly elevated recombination rate, and thus strength of gene conversion relative to background levels, for example, in humans mean *B* is estimated at ~ 0.4 (Glémin *et al.*, 2015), while inside recombination hotspots it reaches as high as ~ 18 (Glémin *et al.*, 2015). In birds, the combination of a karyotype consisting of a few long macro-chromosomes and many smaller micro-chromosomes (Hansson *et al.*, 2010; Stapley *et al.*, 2008; van Oers *et al.*, 2014; Zhang *et al.*, 2014) and obligate crossing over causes large chromosomal differences in recombination rate (Backström *et al.*, 2010; Stapley *et al.*, 2008; van Oers *et al.*, 2014). Additionally, it has been suggested that birds’ lack of PRDM9, has resulted in stable recombination hotspots and conserved recombination characteristics between species (Singhal *et al.*, 2015). Together this is suggested to allow strong gBGC to act on the same region of the genome over a longer time period than in mammals (Rousselle *et al.*, 2019; Singhal *et al.*, 2015), driving GC content increases, with studies reporting that GC content is below GC* content in most avian lineages (Bolívar *et al.*, 2016; Mugal *et al.*, 2013; Rousselle *et al.*, 2019; Weber *et al.*, 2014). Furthermore, some organisms, such as the honey bee *Apis mellifera*, lack pronounced recombination hotspots, yet have very high genome-wide recombination rate with 5 crossovers per arm and correspondingly elevated mean *B* estimates of ~ 5 (Wallberg *et al.*, 2015). Overall, gBGC is seemingly an ubiquitous force with mean *B* estimates varying from 0.4 to 5 across the tree of life (Long *et al.*, 2018).

As *B* is defined as 4*N_e_b*, not only is its strength modulated by recombination rate increasing *b* (the strength of conversion) as outlined above but also by the effective population size (*N_e_*). As such species with larger *N_e_* should have larger *B* and a reduced confounding impact of genetic drift. This has been reported in a few studies, with correlations between *N_e_* and GC content at 3rd codon positions (GC3) in birds, largely driven by increased GC in smaller bodied, larger *N_e_* species, as well as correlations between *N_e_* and GC* (Weber *et al.*, 2014). More recently *B* at fourfold degenerate sites (4-fold sites) has been shown to correlate with *N_e_* in great apes (Borges *et al.*, 2019). However, an analysis of *B* more broadly across animal taxa, failed to yield a relationship with *N_e_* (Galtier *et al.*, 2018). Furthermore, to date the role of *N_e_* is a less well empirically studied aspect of gBGC and little work has looked at fine scale variation in the strength of gBGC between species of differing *N_e_*.

The avian system has been the model of choice for many studies addressing GC evolution and biased gene conversion (Bolívar *et al.*, 2016, 2018, 2019; Corcoran *et al.*, 2017; Gossmann *et al.*, 2018; Rousselle *et al.*, 2019; Weber *et al.*, 2014). The suitability of avian genomes for addressing these topics stems from their variable intra genomic recombination landscapes (Backström *et al.*, 2010; Stapley *et al.*, 2008; van Oers *et al.*, 2014) and conserved recombination hotspots (Singhal *et al.*, 2015) providing a natural experiment for addressing the role of recombination and *N_e_* in gBGC and GC content evolution. In addition, birds’ conserved karyotype and synteny (Hansson *et al.*, 2010; Stapley *et al.*, 2008; van Oers *et al.*, 2014; Zhang *et al.*, 2014) aids between species comparisons.

Of the work on gBGC to date, much has focused on exploring its impact and interaction within genes and coding regions, largely addressing how it confounds signatures of selection (Bolívar *et al.*, 2019; Corcoran *et al.*, 2017; Gossmann *et al.*, 2018; Ratnakumar *et al.*, 2010; Rousselle *et al.*, 2019). Of those studies that have considered the action of gene conversion in the non-coding genome (Duret and Arndt, 2008; Glémin *et al.*, 2015; Haddrill and Charlesworth, 2008; Jackson *et al.*, 2017; Muyle *et al.*, 2011; Wallberg *et al.*, 2015), little work has investigated fine scale variation within the genome and how this compares between species. Here we investigate variation in the strength of gBGC within the non-coding genomes of two passerines, the great tit (*Parus major*) and the zebra finch (*Taeniopygia guttata*), using previously published whole genome resequencing data (Corcoran *et al.*, 2017; Singhal *et al.*, 2015). We seek to address how conserved the gBGC landscape is between these species and how the strength of gBGC has been modulated by the recombination rate and *N_e_* within and between the species.

## Materials and methods

### The dataset

The dataset consisted of 10 European great tits from across the sampling locations in Laine *et al.* (2016), sequenced to a mean coverage of 44X in Corcoran *et al.* (2017) and 10 zebra finches sequenced to a mean coverage of 22X, a subset of individuals from the Fowlers Gap population in Australia from the dataset published in Singhal *et al.* (2015). The dataset is as described in Corcoran *et al.* (2017), but for clarity we will reiterate the main calling pipeline here.

SNP calling was performed using GATK v3.4 (Van der Auwera *et al.*, 2013). Raw genotypes were initially called using the GenotypeGVCF and HaplotypeCaller tools and hard filtered according to the GATK best practice (Van der Auwera *et al.*, 2013). This call set was used as a training set to perform base quality score recalibration (BQSR). Variants were then recalled from the recalibrated BAM files both with GATK as above and also using Freebayes v1.02 (Garrison and Marth, 2012). The intersection of the programs’ calls was taken and SNPs with less than half, or more than double the mean depth, and SNPs with a QUAL score less than 20 were removed. This filtered intersection of SNPs was used as a training set to perform variant quality score recalibration (VQSR) on the GATK called variants. Tranche level thresholds were set at 99% for the zebra finch and 99.9% for the great tit. For both species we obtained VCF files for SNPs and monomorphic sites from Corcoran *et al.* (2017).

Additionally, a three species whole genome alignment between zebra finch (v3.2.4; Warren *et al.*, 2010), great tit (v1.0.4; Laine *et al.*, 2016) and collared flycatcher (*Ficedula albicollis*) (v1.5; Ellegren *et al.*, 2012) was obtained from Barton and Zeng (2019), and a three species alignment between chicken (*Gallus gallus*) (v5.0; Hillier *et al.*, 2004), zebra finch and great tit from Corcoran *et al.* (2017). The former alignment was used to infer the ancestral states of SNPs, and the latter, with the more distant chicken out-group was used to infer substitution rates and ancestral base composition (described later). Both of these alignments were generated as follows. Firstly pairwise alignments were generated with LASTZ (Harris, 2007) between each species and the zebra finch genome, which was used as reference. These alignments were then chained and netted with axtChain and chainNet respectively (Kent *et al.*, 2003). Single coverage was ensured for the zebra finch reference genome using single cov2.v11 from the MULTIZ package, and multiple alignments were created from the pairwise alignments using MULTIZ (Blanchette *et al.*, 2004).

### Annotation and filtering

We assigned the ancestral states for the SNPs using the whole genome alignment (with collared flycatcher) and parsimony based approach, where for each species either the reference allele or the alternate allele had to supported by both out-groups to be assigned as ancestral.

We downloaded the great tit genome annotation (version 1.03) from ftp://ftp.ncbi.nlm.nih.gov/genomes/all/GCF/001/522/545/GCF_001522545.1_Parus_major1.0.3/GCF_001522545.1_Parus_major1.0.3_genomic.gff.gz (last accessed 05/03/19) and the zebra finch annotation from ftp://ftp.ncbi.nlm.nih.gov/genomes/all/GCF/000/151/805/GCF_000151805.1_Taeniopygia_guttata-3.2.4 (last accessed on 05/03/19). We used the annotations to remove variants falling within exons. Additionally coordinates for ultra-conserved non-coding elements (UCNEs) in the zebra finch genome (taeGut1) were obtained from ftp://ccg.vital-it.ch/UCNEbase/custom_tracks_UCSC/UCNEs_taeGut1.bed (last accessed 05/03/19). We identified the corresponding positions in the great tit in the whole genome alignment, before removing any variants falling within UCNEs. Additionally we restricted our analysis to the autosomes, removing the Z chromosome. This left non-coding datasets of putatively neutral variants, numbering 9,800,315 SNPs for great tit, and 29,973,954 SNPs for zebra finch.

From our non-coding SNP dataset we generated an additional subset, with CpG sites excluded, where a CpG site was defined as any site where at least one of the alleles of the site was in a 5’ → 3’ CpG dinucleotide or in a 3’ → 5’ GpC dinucleotide.

### Orthologous window preparation

The zebra finch genome was divided into 1Mb non-overlapping windows and we used the three species whole genome alignment (zebra finch, great tit, collared flycatcher) to identify the aligned sequence and coordinates in the great tit genome and extracted variants and numbers of callable sites from our VCF files. For each window in each species we calculated the GC content using the respective reference genomes. GC content was calculated for all sites in the window, and for non-CpG sites. Secondly, we calculated crossover rate for each window, using the available linkage map data for each species (Stapley *et al.*, 2008; van Oers *et al.*, 2014, for zebra finch and great tit respectively) and the pipeline outlined in Corcoran *et al.* (2017).

### Estimating the strength of gene conversion

We extracted the number of callable sites for weak bases (A and T nucleotides) and strong bases (G and C nucleotides) along with the site frequency spectra for weak to strong mutations (*WS*), strong to weak mutations (*SW*) and weak to weak and strong to strong mutations (*WWSS*) in all windows and datasets. We then applied the *M* 1⁎ model of Glémin *et al.* (2015), implemented in the anavar package (Barton and Zeng, 2018), to all windows with at least 1, 000 SNPs. Briefly, the model estimates the population scaled mutation rate (*θ* = 4*N_e_μ*), the population scaled strength of gBGC (*B* = 4*N_e_b*) and estimates and controls for polarisation error for both *SW* and *WS* mutations using *WW SS* sites as a neutral reference unaffected by gBGC. Demography is controlled for using the method of *Eyre-Walker et al.* (2006), which has been shown previously to obtain similar results to a method that explicitly model recent changes in population size (Jackson *et al.*, 2017).

We performed multiple regressions in R (R Core Team, 2015) to estimate the relative contributions of crossover rate and local *N_e_* (using nucleotide diversity [*π*] as a measure of *N_e_*) in determining *B*, we ran these analysis using crossover rate and separately, GC content as measures of recombination rate. We estimated the relative importance of the predictors (as a proportion of the total variance explained) using the ‘pmvd’ method implemented in the relaimpo package (Groemping, 2006).

### Equilibrium GC content

We estimated the ancestral GC content per window for the lineage leading to great tits and zebra finches using the whole genome alignment (containing chicken, zebra finch and great tit) and the GTR-NH_*b*_ model in baseml within PAML (Yang, 2007). The model allows for non-stationary base content and for independent substitution rates on each branch. From the model we obtained the posterior probabilities of the ancestral states and weighted each ancestral nucleotide by this probability (as in Matsumoto *et al.*, 2015) to reconstruct ancestral GC content with uncertainty incorporated. We then estimated the rate of *WS* substitutions

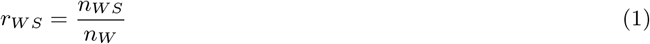

where *n_WS_* is the number of *WS* substitutions and *n_W_* is the number of weak bases (As and Ts) in the ancestral sequence. Similarly we estimated the rate of *SW* substitutions

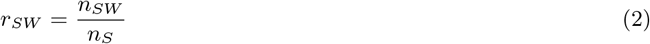

where *n_SW_* is the number of *SW* substitutions and *n_S_* is the number of strong (Gs and Cs) bases in the ancestral sequence. Finally we estimated the equilibrium GC content

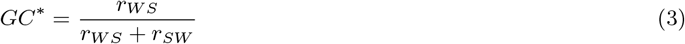

The GTR-NH_*b*_ model was a better fit then the GTR model, which assumes base composition is at equilibrium, for all but five windows as judged by likelihood ratio tests (data not shown). Additionally, the model estimates of 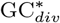 correlated strongly with those derived from parsimony estimates of the substitution rates for both great tit (Pearson’s *r* = 0.94, *p* < 2.2 × 10^−16^) and zebra finch (Pearson’s *r* = 0.96, *p* < 2.2 × 10^−16^), although the mean 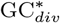 was lower for the model estimates than the parsimony estimates in both species (0.39 versus 0.43 respectively for great tit and 0.38 versus 0.42 respectively for zebra finch).

To obtain a more recent view of the base composition evolution and gBGC we also calculated 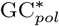 from our application of the Glémin *et al.* (2015) model to our polymorphism dataset. In order to do so we took the estimates of *B* (*B* = 4*N_e_b*) and mutation rates (*θ* = 4*N_e_μ*) estimated per window by anavar and substituted them into

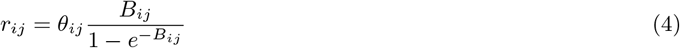

where *r_ij_* is the fixation rate of mutations from *i* to *j* and where *B_WS_* = −*B_SW_* = *B*. The resulting fixation rates were then substituted into equation 3 to obtain GC*.

### Demographic analysis

To investigate the demographic history in the zebra finch and the great tit we fitted demographic models to the data using the VarNe package Zeng *et al.* (2019). The package performs maximum likelihood estimation of a number of population genetic parameters, including *θ* (4*N_e_μ*), the magnitude of a population size change (*g*), the timing of the event (*τ*, in units of 2*N_e_*) and the rate of ancestral state misidentification (*E*), allowing population size changes between a specified number of time points, or epochs, from the site frequency spectrum of a target locus. We applied 1 epoch and 2 epoch models to the summed site frequency spectra for *WWSS* (GC conservative) non-coding SNPs from our window dataset. We tested whether the 2 epoch model (variable population size) was a better fit than the 1 epoch model (constant size), using likelihood ratio tests. We performed 100 rounds of bootstrapping by resampling windows from our window dataset with replacement.

We also applied the 2 epoch model above individually to each window in our dataset to obtain local estimates of the magnitude of *N_e_* change. For these analyses we required windows to have a minimum of 1000 SNPs and windows that failed to return reliable parameter estimates were excluded (67 windows in the great tit, 4 windows in the zebra finch).

In order to infer how much our polymorphism based estimate of the equilibrium GC content 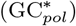 might differ prior to the inferred population size change in each species, we divided our estimates of *B_WS_*, *θ_WS_*, *B_SW_* and *θ_SW_* by a correction factor *C*, as a function of *g* and *τ* estimates per window:

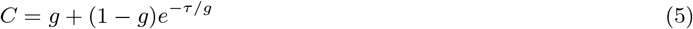

We then substituted the rescaled values into equation 4, to calculate the fixation probabilities for *WS* and *SW* polymorphisms under the reduced *B* scenario. The fixation probabilities were then substituted into equation 3 to calculate *GC*\*.

### Data availability

All scripts and command lines used in the analysis pipeline can be found at: https://github.com/henryjuho/biased_gene_conversion. The VCF files, whole genome alignments and orthologous window coordinates are accessible at: **link**.

## Results

### Summary of the window dataset

We used a whole genome alignment between zebra finch, great tit and collared fly catcher (*Ficedula albicollis*) to identify 1Mb orthologous windows between the zebra finch and great tit. This resulted in 904 1Mb windows in zebra finch genome and 898 orthologous windows in the great tit genome (table 1). The lower number of great tit windows is due to gaps in the whole genome alignment. We used the respective genome annotations to identify non-coding regions within these windows, in which we identified single nucleotide polymorphisms (SNPs) using a resequencing dataset of 10 zebra finches (from Singhal *et al.*, 2015) and 10 great tits (from Corcoran *et al.*, 2017). This resulted in similar numbers of callable sites in both species, roughly 500,000 bp per 1 Mb window; this drop is a result of our focus on non-coding regions (excluding ultra-conserved non-coding elements [UCNEs]), and our maximum parsimony approach to assigning ancestral states, which is dependant on coverage of all species in our whole genome alignment and no ambiguity between out-groups. When considering variants per window, we see that the mean number of variants is higher in the zebra finch, consistent with a larger effective population size in the zebra finch (Corcoran *et al.*, 2017). We see very similar mean GC content and mean crossover rates in both species, with strong correlations between the two species’ GC content (Pearson’s *r* = 0.83, *p* = 1.6 × 10^−230^, figure S1a) and crossover rate (Spearman’s *ρ* = 0.72, *p* = 2.6 × 10^−140^, figure S1b) across the dataset, as well as positive correlations between GC content and crossover rate within each species (great tit: Spearman’s *ρ* = 0.57, *p* = 3.8 × 10^−79^, zebra finch: Spearman’s *ρ* = 0.53, *p* = 4.2 × 10^−67^, figure S2).

**Table 1:**
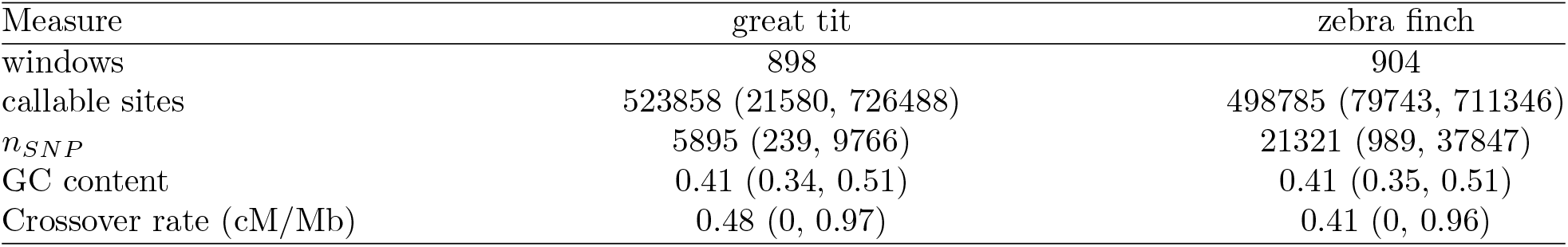
Summary of the window dataset, showing means and the 2.5 and 97.5 percentiles in brackets. Crossover rates are log10 transformed.

| Measure | great tit | zebra finch |
| --- | --- | --- |
| windows | 898 | 904 |
| callable sites | 523858 (21580, 726488) | 498785 (79743, 711346) |
| $n_{SNP}$ | 5895 (239, 9766) | 21321 (989, 37847) |
| GC content | 0.41 (0.34, 0.51) | 0.41 (0.35, 0.51) |
| Crossover rate (cM/Mb) | 0.48 (0, 0.97) | 0.41 (0, 0.96) |

### The strength of gene conversion correlates with crossover rate and *N_e_*

To estimate the population scaled strength of gBGC (*B*), we applied the Glémin *et al.* (2015) model to each window in our dataset. The resulting estimates of *B* positively correlate with both crossover rate and *π* (as a proxy for local *N_e_*, allowing us to separate the contributions of *N_e_* to the compound parameter *B* = 4*N_e_b*) in both the great tit and the zebra finch (table 2, figure 1). The relationships are stronger when using mean GC content as a proxy for recombination rate in both species (table 2, figure S3) and all relationships are maintained when performed on a dataset filtered for CpG sites (table S1). Crossover rate or mean GC content explains a larger proportion of the total variance (80 − 95%) than *π* within both species (table 2, table S1).

**Table 2:**
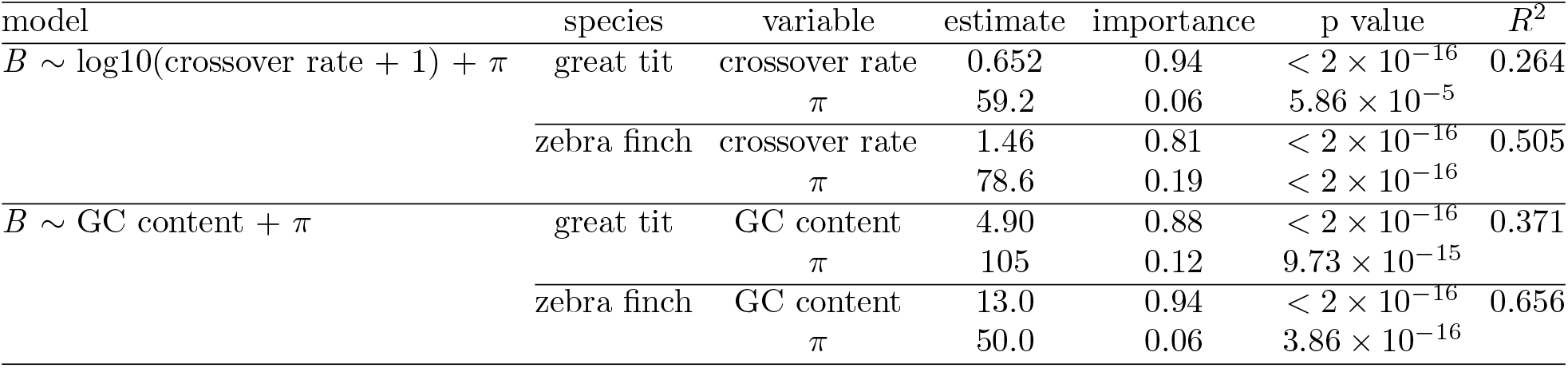
Results of multiple regression analysis of the strength of gene conversion (*B*) against GC content and *π*, and against crossover rate and *π*, separately, for both species. Importance is the relative importance (as a proportion of the total variance explained) as estimated using the pmvd method implemented in the `relaimpo` package (Groemping, 2006).

**Figure 1:**
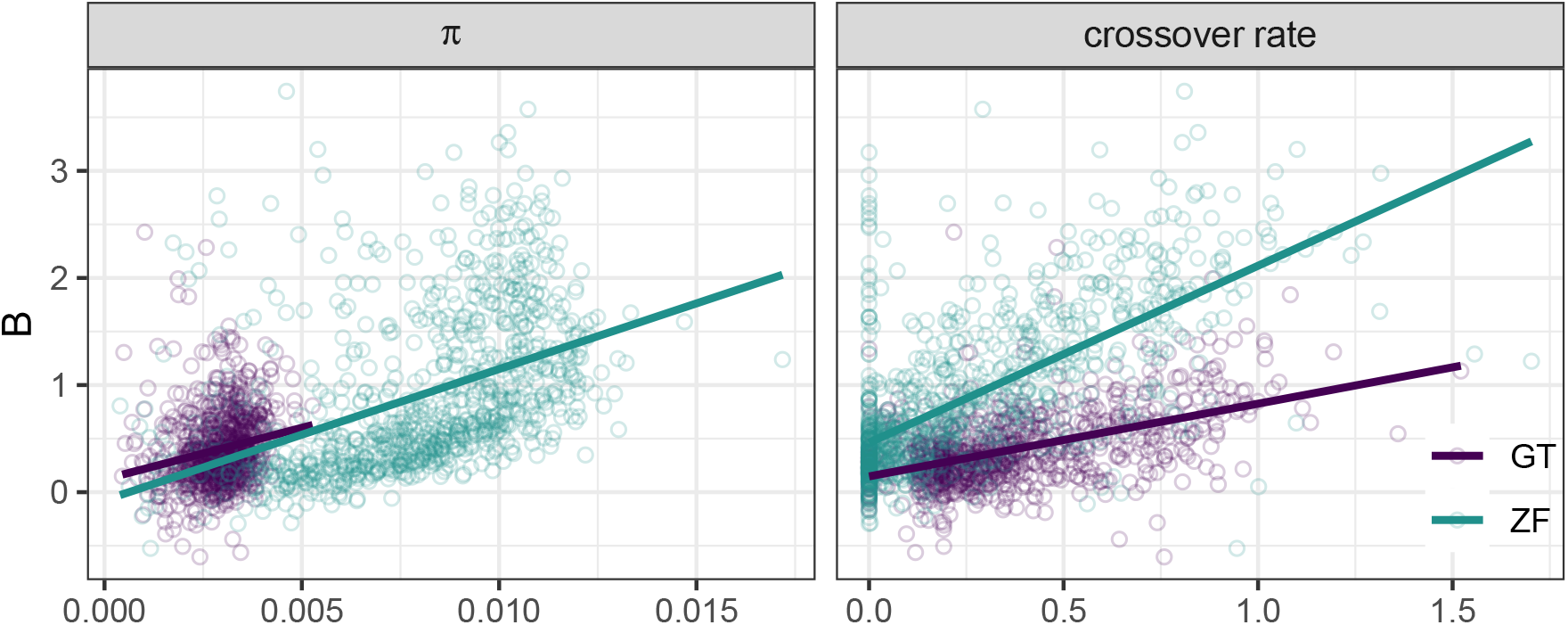
The relationship between nucleotide diversity (*π*) and the strength of gene conversion (*B*) (left panel) and mean window crossover rate and *B* (right panel) in the great tit (purple) and zebra finch (turquoise). Multiple regression results can be seen in table 2.

### *B* is correlated between the species

Comparison of the model estimates of *B* between zebra finch and great tit show a significantly larger mean *B* value in zebra finch 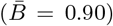 than great tit 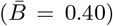 (Wilcoxon rank sum, *W* = 491903, *p* = 2.5 × 10^−49^; figure 2a), inline with the species’ twofold difference in *N_e_* (Corcoran *et al.*, 2017). However, when we standardise our *B* estimates by *π* as a measure of *N_e_*, the difference between the two species is greatly reduced and the distributions of *B/π* are similar in both species (figure 2b). However, *B/π* is slightly, but significantly larger in the great tit (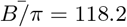 and 80.8 for great tit and zebra finch respectively, Wilcoxon rank sum, *W* = 305880, *p* = 6.1 × 10^−10^). We also see a positive correlation between the ratio of the species’ nucleotide diversity (*π_zf_ /π_gt_*) and the ratio of the species’ *B* (*B_zf_ /B_gt_*) (Spearman’s *ρ* = 0.44, *p* < 2.2 × 10^−16^), supporting the idea that *N_e_* drives the between species differences in *B*. Furthermore, we see a strong correlation between *B* in the great tit and *B* in the zebra finch (Pearson’s *r* = 0.50, *p* < 2.2 × 10^−16^, figure 3) as well as between *B/π* in great tit and *B/π* in zebra finch (Pearson’s *r* = 0.38, *p* < 2.2 × 10^−16^), in keeping with the conserved crossover rate and GC content between species reported above.

**Figure 2:**
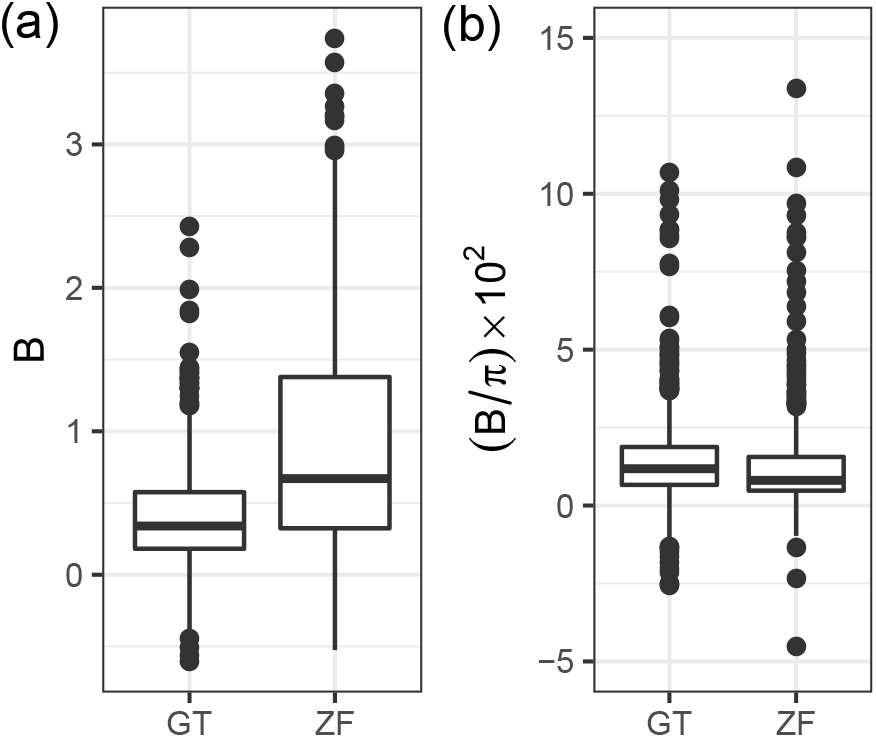
Comparison of the distribution of *B* values (population scaled strength of biased gene conversion) (a) and *B* standardised by *π* as a proxy for the effective population size *N_e_* (b) between the great tit (GT) and zebra finch (ZF). The y axis for b has been cropped for clarity.

**Figure 3:**
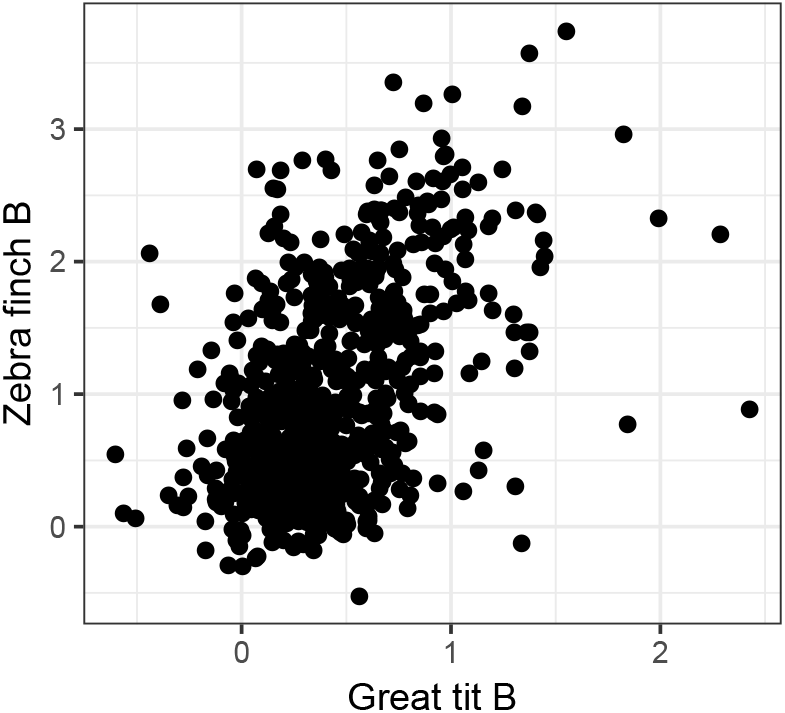
The strength of biased gene conversion (*B*) in the zebra finch positively correlates with *B* in the great tit.

### Equilibrium GC content

To assess the longer term GC dynamics of both the great tit and zebra finch genomes, we calculated the equilibrium GC content (GC*), which is the GC content that when reached will result in equal numbers of GC alleles fixed as lost.

Firstly, we calculated GC* using divergence data 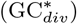 for each lineage, using the WS and SW substitution rates estimated in PAML (see methods). This provides a long term average of GC* since the two species diverged. This gave a mean 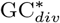 of 0.39 for great tit and 0.38 for zebra finch, both of which are similar to, but significantly below, the mean GC contents in our alignment datasets of 0.40 for both great tit (Wilcoxon rank sum, *W* = 282790, *p* = 1.1 × 10^−8^) and zebra finch (Wilcoxon rank sum, *W* = 241190, *p* < 2.2 × 10^−16^) (figure 4). Note the alignment dataset is a subset of the main dataset (as coverage is required across all species in the chicken/zebra finch/great tit alignment) and yields slightly lower mean GC than reported in table 1. *B* positively correlates with 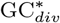 in both great tit (Pearson’s *r* = 0.54, *p* < 2.28 × 10^−55^) and zebra finch (Pearson’s *r* = 0.81, *p* < 8.22 × 10^−181^). Similar relationships are seen between 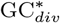 and crossover rate (Spearman’s *ρ* = 0.55, *p* = 6.02 × 10^−62^ for great tit and Spearman’s *ρ* = 0.66, *p* = 3.85 × 10^−98^ for zebra finch) and between 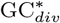 and current GC content (Pearson’s *r* = 0.56, *p* = 9.32 × 10^−65^ for great tit and Pearson’s *r* = 0.77, *p* = 1.49 × 10^−148^ for zebra finch).

**Figure 4:**
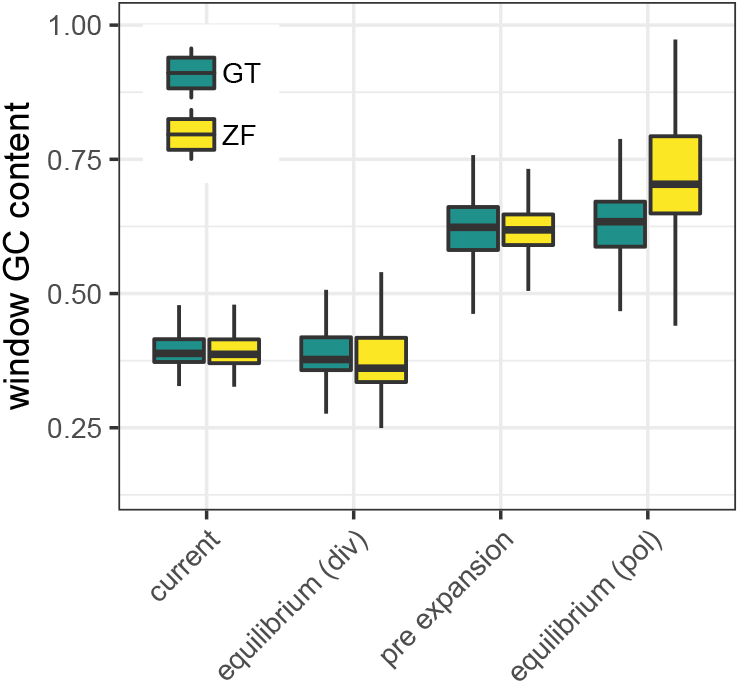
Per window estimates of current GC content, equilibrium GC content from both divergence data (div) and polymorphism data (pol) and estimates of equilibrium GC content before expansion (pre expansion) for both species.

Secondly, to look at base composition evolution over a more recent time scale we also calculated GC* from polymorphism data, using our *θ* and *B* estimates derived from the Glémin *et al.* (2015) model (see methods), henceforth 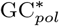. This approach yielded markedly higher equilibrium GC content estimates than the substitution rate based approach, for both great tit (Wilcoxon rank sum, *W* = 518421, *p* = 1.48 × 10^−225^,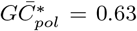) and zebra finch (Wilcoxon rank sum, *W* = 575196, *p* = 1.24 × 10^−245^,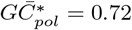)

### Evidence of population expansions

In order to understand the effects of recent demographic changes on the difference between our longer term measures of 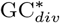 and our more recent 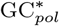 estimates, we fitted demographic models to each species using the VarNe package (Zeng *et al.*, 2019). The models estimate the magnitude (*g*) and timing of population size changes (*τ*, in units of 2*N_e_*) between different time points or ‘epochs’. In both the zebra finch and the great tit a 2 epoch model (table S3) fit the data significantly better than a 1 epoch model (i.e. a model with constant population size) as judged by likelihood ratio tests. For the zebra finch we estimate a *g* of 12.3 and *τ* of 1.25, suggesting a large population expansion ~ 495 thousand years ago (table S3). In the great tit we see lower values with a *g* of 1.89 and *τ* of 0.208, characterising a smaller, more recent population expansion ~ 140 thousand years ago (table S3).

### Local *N_e_* increase correlates with increases in equilibrium GC content in the zebra finch

Nucleotide diversity is positively correlated with recombination rate in both the great tit and zebra finch (Corcoran *et al.*, 2017), showing *N_e_* varies locally within their genomes. As loci with differing *N_e_* can respond differently to a shared demographic change (see Zeng *et al.*, 2019), we sort to investigate how historical changes in local *N_e_* have impacted equilibrium GC content, and the difference between our 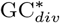 and 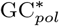 estimates. In each species, we refitted the ‘2 epoch’ model in `VarNe`, to each window in our orthologous window dataset. The mean maximum likelihood parameter estimates across all windows agreed with those from the model fitted to the dataset as a whole, although were slightly higher, probably a result of our requirement of a minimum of 1000 SNPs per window to provide sufficient power, excluding the lowest *N_e_* windows (table S4).

For each window, we divided our estimates of *θ* and *B* from the Glémin *et al.* (2015) model by a rescaling factor *C* (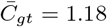 and 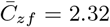), a function of the window’s *g* and *τ* estimates (see equation 5) to control for the effects of recent population expansion. We used these rescaled values to obtain per window estimates of GC*, prior to the inferred local *N_e_* increases. This approach yielded a mean pre-expansion GC* of 0.62 in both the zebra finch and the great tit (figure 4), demonstrating the transient effect of recent population size changes on equilibrium GC content. These GC* estimates are still high relative to 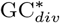, potentially due to population expansions taking place before the most recent common ancestor of the polymorphism samples.

Additionally, we compared the per window values of *C* (a measure of the impact of *N_e_* increase on *B*) with the difference between our two GC* estimates. This returned a significant positive correlation between GC* increase 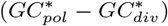 and *C* in the zebra finch (Spearman’s *ρ* = 0.46, *p* < 2.2 × 10^−16^) and a weak positive correlation in the great tit (Spearman’s *ρ* = 0.081, *p* = 0.036). The stronger correlation in the zebra finch is consistent with an older and larger expansion in this species providing more time for evolution to influence *C* and GC*.

## Discussion

Most contemporary studies on the role of GC biased gene conversion (gBGC) in genome evolution have focused on coding regions where gBGC is confounded by selection, (Bolívar *et al.*, 2019; Corcoran *et al.*, 2017; Gossmann *et al.*, 2018; Ratnakumar *et al.*, 2010; Rousselle *et al.*, 2019) and processes like codon usage bias (Haddrill *et al.*, 2008; Jackson *et al.*, 2017). Additionally few of these studies have looked at the impact of *N_e_* on the strength of gBGC. Here we analyse re-sequencing data for 10 great tits (Corcoran *et al.*, 2017), and 10 zebra finches (Singhal *et al.*, 2015). Using non-overlapping 1Mb orthologous windows, we investigate how the strength and impact of gBGC varies both within and between the non-coding genomes of these birds.

### The strength of gene conversion is modulated by *N_e_*

Our mean estimates of *B* in the great tit and the zebra finch of 0.40 and 0.90 respectively, are similar to mean genome wide estimates of *B* in humans of 0.38 (Glémin *et al.*, 2015), and fall at the lower end of the *B* range of 0.4 to 5 reported by Long *et al.* (2018) in a comparative study with taxa from across the tree of life. Mutations with *N_e_s* < 1 (here *B* = 4*N_e_b* < 4) are considered effectively neutral, our mean *B* estimates fall below 1, suggesting gBGC in the non-coding regions of these species is operating at low efficiency.

*B* correlates with both recombination rate and *π* in these species (table 2) suggesting both parameters are modulating *B* in their genomes, although recombination rate has the larger impact (when measured by crossover rate or mean GC content), particularly in the great tit. This is consistent with elevated gBGC in regions with higher recombination rate in humans (Glémin *et al.*, 2015) and correlations between GC content at 4-fold sites and recombination rate in flycatchers (Bolívar *et al.*, 2016), although these analyses did not control for local *N_e_*. When using GC content as a measure of recombination rate instead of crossover rate these relationships are strengthened. This may reflect that GC content is a better measure of long term recombination rate, that our crossover rate estimates are constrained by the density of the linkage maps available (Stapley *et al.*, 2008; van Oers *et al.*, 2014), lower variance in our GC estimates (table 1), or a mixture of the three.

The conservation of the biased gene conversion landscape between the zebra finch and great tit, as seen by the strong correlation of window *B* estimates between the species, is relatively intuitive with GC content and crossover rate also correlating well between the species and likely a result of birds’ conserved recombination hotspots (Singhal *et al.*, 2015), karyotype and synteny (Hansson *et al.*, 2010; Stapley *et al.*, 2008; van Oers *et al.*, 2014; Zhang *et al.*, 2014). Consistently, we also see similar mean crossover rates in each species (table 1).

Nonetheless, mean *B* is approximately twofold higher in the zebra finch. As *B* is the product of *b* (the strength of biased gene conversion) and *N_e_*, either parameter could be driving this increase. When we standardise *B* by *π* (as a measure of *N_e_*), the between species difference is greatly reduced. This, combined with the correlation of the ratios of between species *B* (*B_zf_ /B_gt_*) and *π* (*π_zf_ /π_gt_*), suggests the twofold larger *N_e_* in the zebra finch (Corcoran *et al.*, 2017) is elevating its *B*. This also implies that *b* is comparable between the species and has remained relatively stable since their divergence. Consistently, GC3 content correlates with *N_e_* (using life history traits as proxies) across the avian phylogeny (Weber *et al.*, 2014). More broadly, it fits with findings in great apes, where *B* at 4-fold sites correlates with *N_e_* (Borges *et al.*, 2019) and amongst rice species (*Oryza spp.*), where selfing species (with reduced *N_e_*) also have lower *B* estimates (Muyle *et al.*, 2011). However, a recent analysis by Galtier *et al.* (2018) between more diverged species failed to find a relationship between *B* and *N_e_*, with the authors suggesting that *b* may be inversely related to *N_e_* between distant taxa, and only remain homogenous within groups, such as birds, suggesting that *B* only responds to *N_e_* over small time-scales.

### Non-coding equilibrium GC content

Our two measures of equilibrium GC content (GC*, the theoretical GC content at which the same number of GC alleles are fixed as AT alleles, and thus stable GC content is reached), from divergence data 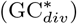 and polymorphism data 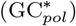, respectively provide a longer term and more recent insight into GC*.

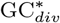 is similar to, albeit significantly lower than, current GC content in both species. This is at odds with previous avian studies where GC content is below GC* in most lineages (Bolívar *et al.*, 2016; Rousselle *et al.*, 2019; Weber *et al.*, 2014). However, these studies focus on coding regions, which are have elevated GC content and recombination rates over non-coding regions in birds (Singhal *et al.*, 2015; Weber *et al.*, 2014); in our dataset, GC content is ~ 10%higher in coding regions than non-coding regions (table S2). Consequently, gBGC is likely stronger in coding regions, as suggested by 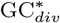 estimates of 0.6 − 0.8 at fourfold sites in collared flycatcher (Bolívar *et al.*, 2016) and a median 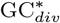 of 0.6 at 3rd codon positions across 48 bird species (Weber *et al.*, 2014), compared to our non-coding 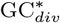 of 0.39 in the great tit and 0.38 in the zebra finch. These differing dynamics may be contributed to by the avian micro-chromosomes which are characterised by high gene density, and high recombination rates stemming from obligate crossing over and their short length (Burt, 2002; Stapley *et al.*, 2008; van Oers *et al.*, 2014). Equally, if codon usage bias (CUB) is operating in addition to gBGC (de Procé *et al.*, 2012; Galtier *et al.*, 2018) and favours G and C ending codons (de Procé *et al.*, 2012) this could elevate avian coding GC over non-coding GC, and also inflate estimates of gBGC in coding regions, however, evidence for CUB in birds is lacking. Overall, it seems these regions have been evolving towards different equilibria, similar to some species of rice (Muyle *et al.*, 2011), with weak gBGC allowing for more AT biased fixation patterns (see McVean and Charlesworth, 1999) and a slightly decreasing GC content in non-coding regions since the great tit zebra finch split.

### The effect of demography on *B* and GC*

Our mean 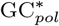 estimates are higher than our 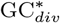 values, 0.63 versus 0.39 for great tit and 0.72 versus 0.38 for zebra finch. 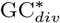 represents a long term average of GC* since the divergence of the great tit and zebra finch lineages 40 to 45 million years ago (Barker *et al.*, 2004), whereas 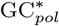 provides a more recent snapshot, of the order of 4*N_e_* generations ago, around ~ 3.5 and ~ 4.3 million years ago for the great tit and zebra finch respectively (estimated using the current *N_e_* estimates and generation times in table S5). Consequently, our higher 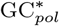 estimates suggests *B* is currently higher than the long term average for the species, this is the opposite to what is seen in *Drosophila melanogaster*, where longer term estimates of *B* are higher than those from the Glémin *et al.* (2015) model (Jackson *et al.*, 2017). As *B* is the product of *b* (the underlying strength of conversion bias) and *N_e_*, this increase could be driven by increases in the population size and/or *b* through changing recombination rates. As recombination rates are relatively stable and conserved in these species (Singhal *et al.*, 2015; van Oers *et al.*, 2014, this study), it seems more probable the current elevation of *B* is driven by changes in *N_e_*.

Here, we estimate ~ 12-fold and ~ 2-fold population expansions for the zebra finch and great tit respectively, in agreement with previous evidence for expansions in both species (Balakrishnan and Edwards, 2008; Corcoran *et al.*, 2017; Laine *et al.*, 2016). The magnitude of the great tit expansion is similar to reported values of 2.75 (Laine *et al.*, 2016), 2.31 (Corcoran *et al.*, 2017) and 1.68 (Hayes *et al.*, 2020). The zebra finch expansion magnitude of 12.3 is close to the estimate of 10 from Corcoran *et al.* (2017), the upper limit of the method used. The larger increase in *N_e_* for the zebra finch is consistent with the greater difference in GC* measures in this species (figure 4). Furthermore, our estimates of 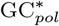 corrected for the inferred population expansions are 0.62 in both species, suggesting each species’ average *N_e_* have remained similar since they diverged. The difference between 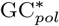 and 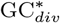 is reduced by 29%after correction in the zebra finch, but only by 4% in the great tit. Concordantly, the difference between 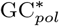 and 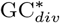 correlates well with our correction factor *C*, a measure of the impact of *N_e_* increase, in zebra finch only. As the polymorphism data spans at most 10% of the species divergence time, most of the demographic history since the their split is not captured in our analysis, thus the modest impact of the recent expansions on GC* is perhaps unsurprising.

## Conclusion

We show that the underlying strength of gene conversion *b* is conserved between the great tit and zebra finch, with the zebra finch’s larger population scaled strength of gBGC, *B*, due to its larger effective population size. Within each species’ genome, variation in *B* is driven by variation in both recombination rate and local *N_e_*, with the former having the larger impact.

When considering the equilibrium GC content, we see that 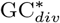 and GC* prior to the inferred population expansions are similar between the great tit and zebra finch, suggesting that they have had similar average *N_e_* since their divergence. Our higher 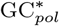 estimates are likely explained by the short timescale covered by the polymorphism data relative to the divergence data.

## Acknowledgements

We thank Pádraic Corcoran for providing an initial implementation of the window pipeline and Brian Charlesworth for his comments on the manuscript. This work was supported by a PhD studentship funded by the Department of Animal and Plant Sciences, University of Sheffield, to H.J.B. Support was also provided by the Natural Environment Research Council via a research grant awarded to K.Z. (NE/L005328/1). The analyses were performed on the University of Sheffield’s high performance computing cluster ‘ShARC’.

## References

Backström, N., Forstmeier, W., Schielzeth, H., Mellenius, H., Nam, K., Bolund, E., Webster, M. T., Ost, T., Schneider, M., Kempenaers, B., and Ellegren, H. 2010. The recombination landscape of the zebra finch Taeniopygia guttata genome. Genome Res, 20(4): 485–95.

Balakrishnan, C. N. and Edwards, S. V. 2008. Nucleotide Variation, Linkage Disequilibrium and Founder-Facilitated Speciation in Wild Populations of the Zebra Finch (Taeniopygia guttata). Genetics, 181(2): 645–660.

Barker, F. K., Cibois, A., Schikler, P., Feinstein, J., and Cracraft, J. 2004. Phylogeny and diversification of the largest avian radiation. Proceedings of the National Academy of Sciences, 101(30): 11040–11045.

Barton, H. J. and Zeng, K. 2018. New Methods for Inferring the Distribution of Fitness Effects for INDELs and SNPs. Molecular Biology and Evolution, 35(6): 1536–1546.

Barton, H. J. and Zeng, K. 2019. The Impact of Natural Selection on Short Insertion and Deletion Variation in the Great Tit Genome. Genome Biology and Evolution, 11(6): 1514–1524.

Baudat, F., Buard, J., Grey, C., Fledel-Alon, A., Ober, C., Przeworski, M., Coop, G., and Massy, B. d. 2010. PRDM9 Is a Major Determinant of Meiotic Recombination Hotspots in Humans and Mice. Science, 327(5967): 836–840.

Blanchette, M., Kent, W. J., Riemer, C., Elnitski, L., Smit, A. F. A., Roskin, K. M., Baertsch, R., Rosenbloom, K., Clawson, H., Green, E. D., Haussler, D., and Miller, W. 2004. Aligning Multiple Genomic Sequences With the Threaded Blockset Aligner. Genome Research, 14(4): 708–715.

Bolívar, P., Mugal, C. F., Nater, A., and Ellegren, H. 2016. Recombination rate variation modulates gene sequence evolution mainly via GC-biased gene conversion, not Hill–Robertson interference, in an avian system. Molecular biology and evolution, 33(1): 216–227.

Bolívar, P., Mugal, C. F., Rossi, M., Nater, A., Wang, M., Dutoit, L., and Ellegren, H. 2018. Biased Inference of Selection Due to GC-Biased Gene Conversion and the Rate of Protein Evolution in Flycatchers When Accounting for It. Molecular Biology and Evolution, 35(10): 2475–2486.

Bolívar, P., Guéguen, L., Duret, L., Ellegren, H., and Mugal, C. F. 2019. GC-biased gene conversion conceals the prediction of the nearly neutral theory in avian genomes. Genome Biology, 20(1): 5.

Borges, R., Szöllősi, G. J., and Kosiol, C. 2019. Quantifying GC-Biased Gene Conversion in Great Ape Genomes Using Polymorphism-Aware Models. Genetics, 212(4): 1321–1336.

Burt, D. W. 2002. Origin and evolution of avian microchromosomes. Cytogenetic and Genome Research, 96(1-4): 97–112.

Chamary, J. and Hurst, L. D. 2005. Evidence for selection on synonymous mutations affecting stability of mRNA secondary structure in mammals. Genome Biology, 6(9): R75.

Chen, J.-M., Cooper, D. N., Chuzhanova, N., Férec, C., and Patrinos, G. P. 2007. Gene conversion: mechanisms, evolution and human disease. Nature Reviews Genetics, 8(10): 762–775.

Comeron, J. M., Ratnappan, R., and Bailin, S. 2012. The Many Landscapes of Recombination in Drosophila melanogaster. PLOS Genetics, 8(10): e1002905. Publisher: Public Library of Science.

Corcoran, P., Gossmann, T. I., Barton, H. J., Great Tit HapMap Consortium, Slate, J., and Zeng, K. 2017. Determinants of the Efficacy of Natural Selection on Coding and Noncoding Variability in Two Passerine Species. Genome Biol Evol, 9(11): 2987–3007.

de Procé, S. M., Zeng, K., Betancourt, A. J., and Charlesworth, B. 2012. Selection on codon usage and base composition in Drosophila americana. Biology Letters, 8(1): 82–85. Publisher: Royal Society.

Duret, L. and Arndt, P. F. 2008. The Impact of Recombination on Nucleotide Substitutions in the Human Genome. PLOS Genetics, 4(5): e1000071.

Duret, L. and Galtier, N. 2009. Biased Gene Conversion and the Evolution of Mammalian Genomic Landscapes. Annual Review of Genomics and Human Genetics, 10(1): 285–311.

Ellegren, H., Smeds, L., Burri, R., Olason, P. I., Backström, N., Kawakami, T., Künstner, A., Mäkinen, H., Nadachowska-Brzyska, K., Qvarnström, A., Uebbing, S., and Wolf, J. B. W. 2012. The genomic landscape of species divergence in Ficedula flycatchers. Nature, 491(7426): 756–760.

Eyre-Walker, A. and Hurst, L. D. 2001. The evolution of isochores. Nature Reviews Genetics, 2(7): 549.

Eyre-Walker, A., Woolfit, M., and Phelps, T. 2006. The distribution of fitness effects of new deleterious amino acid mutations in humans. Genetics, 173(2): 891–900.

Galtier, N. and Duret, L. 2007. Adaptation or biased gene conversion? Extending the null hypothesis of molecular evolution. Trends in Genetics, 23(6): 273–277.

Galtier, N., Roux, C., Rousselle, M., Romiguier, J., Figuet, E., Glémin, S., Bierne, N., and Duret, L. 2018. Codon Usage Bias in Animals: Disentangling the Effects of Natural Selection, Effective Population Size, and GC-Biased Gene Conversion. Molecular Biology and Evolution, 35(5): 1092–1103.

Garrison, E. and Marth, G. 2012. Haplotype-based variant detection from short-read sequencing. arXiv:1207.3907 [q-bio].

Glémin, S., Arndt, P. F., Messer, P. W., Petrov, D., Galtier, N., and Duret, L. 2015. Quantification of GC-biased gene conversion in the human genome. Genome Research, 25(8): 1215–1228.

Gossmann, T. I., Bockwoldt, M., Diringer, L., Schwarz, F., and Schumann, V.-F. 2018. Evidence for Strong Fixation Bias at 4-fold Degenerate Sites Across Genes in the Great Tit Genome. Frontiers in Ecology and Evolution, 6.

Groemping, U. 2006. Relative Importance for Linear Regression in R: The Package relaimpo. Journal of Statistical Software, 17(1): 1–27. Number: 1.

Gutz, H. and Leslie, J. F. 1976. Gene Conversion: A Hitherto Overlooked Parameter in Population Genetics. Genetics, 83(4): 861–866.

Haddrill, P. R. and Charlesworth, B. 2008. Non-neutral processes drive the nucleotide composition of non-coding sequences in Drosophila. Biology Letters, 4(4): 438–441.

Haddrill, P. R., Bachtrog, D., and Andolfatto, P. 2008. Positive and Negative Selection on Noncoding DNA in Drosophila simulans. Molecular Biology and Evolution, 25(9): 1825–1834.

Halligan, D. L. and Keightley, P. D. 2006. Ubiquitous selective constraints in the Drosophila genome revealed by a genome-wide interspecies comparison. Genome Research, 16(7): 875–884.

Hansson, B., Ljungqvist, M., Dawson, D. A., Mueller, J. C., Olano-Marin, J., Ellegren, H., and Nilsson, J.-A. 2010. Avian genome evolution: insights from a linkage map of the blue tit (Cyanistes caeruleus). Heredity, 104(1): 67–78.

Harris, R. S. 2007. Improved pairwise alignment of genomic DNA. Ph.D. Thesis, The Pennsylvania State University.

Hayes, K., Barton, H. J., and Zeng, K. 2020. A study of faster-Z evolution in the great tit (Parus major). Genome Biology and Evolution.

Hillier, L. W., Miller, W., Birney, E., Warren, W., Hardison, R. C., Ponting, C. P., Bork, P., Burt, D. W., Groenen, M. A., Delany, M. E., and others 2004. Sequence and comparative analysis of the chicken genome provide unique perspectives on vertebrate evolution. Nature, 432(7018): 695–716.

Hodgkinson, A. and Eyre-Walker, A. 2011. Variation in the mutation rate across mammalian genomes. Nat. Rev. Genet., 12(11): 756–766.

Hwang, D. G. and Green, P. 2004. Bayesian Markov chain Monte Carlo sequence analysis reveals varying neutral substitution patterns in mammalian evolution. Proceedings of the National Academy of Sciences of the United States of America, 101(39): 13994–14001.

Jackson, B. C., Campos, J. L., Haddrill, P. R., Charlesworth, B., and Zeng, K. 2017. Variation in the Intensity of Selection on Codon Bias over Time Causes Contrasting Patterns of Base Composition Evolution in Drosophila. Genome Biology and Evolution, 9(1): 102–123.

Kent, W. J., Baertsch, R., Hinrichs, A., Miller, W., and Haussler, D. 2003. Evolution’s cauldron: Duplication, deletion, and rearrangement in the mouse and human genomes. Proceedings of the National Academy of Sciences, 100(20): 11484–11489.

Kunstner, A., Nabholz, B., and Ellegren, H. 2011. Significant Selective Constraint at 4-Fold Degenerate Sites in the Avian Genome and Its Consequence for Detection of Positive Selection. Genome Biology and Evolution, 3(0): 1381–1389.

Laine, V. N., Gossmann, T. I., Schachtschneider, K. M., Garroway, C. J., Madsen, O., Verhoeven, K. J. F., de Jager, V., Megens, H.-J., Warren, W. C., Minx, P., Crooijmans, R. P. M. A., Corcoran, P., Great Tit HapMap Consortium, Sheldon, B. C., Slate, J., Zeng, K., van Oers, K., Visser, M. E., and Groenen, M. A. M. 2016. Evolutionary signals of selection on cognition from the great tit genome and methylome. Nat Commun, 7: 10474.

Liu, H., Huang, J., Sun, X., Li, J., Hu, Y., Yu, L., Liti, G., Tian, D., Hurst, L. D., and Yang, S. 2018. Tetrad analysis in plants and fungi finds large differences in gene conversion rates but no GC bias. Nature Ecology & Evolution, 2(1): 164–173.

Long, H., Sung, W., Kucukyildirim, S., Williams, E., Miller, S. F., Guo, W., Patterson, C., Gregory, C., Strauss, C., Stone, C., Berne, C., Kysela, D., Shoemaker, W. R., Muscarella, M. E., Luo, H., Lennon, J. T., Brun, Y. V., and Lynch, M. 2018. Evolutionary determinants of genome-wide nucleotide composition. Nature Ecology & Evolution, 2(2): 237–240.

Matsumoto, T., Akashi, H., and Yang, Z. 2015. Evaluation of Ancestral Sequence Reconstruction Methods to Infer Nonstationary Patterns of Nucleotide Substitution. Genetics, 200(3): 873–890.

McVean, G. a. T. and Charlesworth, B. 1999. A population genetic model for the evolution of synonymous codon usage: patterns and predictions. Genetics Research, 74(2): 145–158.

Mugal, C. F., Arndt, P. F., and Ellegren, H. 2013. Twisted signatures of GC-biased gene conversion embedded in an evolutionary stable karyotype. Molecular Biology and Evolution, 30(7): 1700–1712.

Muyle, A., Serres-Giardi, L., Ressayre, A., Escobar, J., and Glémin, S. 2011. GC-Biased Gene Conversion and Selection Affect GC Content in the Oryza Genus (rice). Molecular Biology and Evolution, 28(9): 2695–2706.

Nagylaki, T. 1983. Evolution of a finite population under gene conversion. Proceedings of the National Academy of Sciences, 80(20): 6278–6281.

Parvanov, E. D., Petkov, P. M., and Paigen, K. 2010. Prdm9 Controls Activation of Mammalian Recombination Hotspots. Science, 327(5967): 835–835.

R Core Team 2015. R: A Language and Environment for Statistical Computing. R Foundation for Statistical Computing, Vienna, Austria.

Rajic, Z. A., Jankovic, G. M., Vidovic, A., Milic, N. M., Skoric, D., Pavlovic, M., and Lazarevic, V. 2005. Size of the protein-coding genome and rate of molecular evolution. Journal of Human Genetics, 50(5): 217–229.

Ratnakumar, A., Mousset, S., Glémin, S., Berglund, J., Galtier, N., Duret, L., and Webster, M. T. 2010. Detecting positive selection within genomes: the problem of biased gene conversion. Philosophical Transactions of the Royal Society B: Biological Sciences, 365(1552): 2571–2580.

Rousselle, M., Laverré, A., Figuet, E., Nabholz, B., and Galtier, N. 2019. Influence of Recombination and GC-biased Gene Conversion on the Adaptive and Nonadaptive Substitution Rate in Mammals versus Birds. Molecular Biology and Evolution, 36(3): 458–471.

Ségurel, L., Wyman, M. J., and Przeworski, M. 2014. Determinants of Mutation Rate Variation in the Human Germline. Annual Review of Genomics and Human Genetics, 15(1): 47–70.

Singhal, S., Leffler, E. M., Sannareddy, K., Turner, I., Venn, O., Hooper, D. M., Strand, A. I., Li, Q., Raney, B., Balakrishnan, C. N., Griffith, S. C., McVean, G., and Przeworski, M. 2015. Stable recombination hotspots in birds. Science, 350(6263): 928–32.

Smeds, L., Mugal, C. F., Qvarnström, A., and Ellegren, H. 2016. High-Resolution Mapping of Crossover and Non-crossover Recombination Events by Whole-Genome Re-sequencing of an Avian Pedigree. PLOS Genetics, 12(5): e1006044.

Stapley, J., Birkhead, T. R., Burke, T., and Slate, J. 2008. A Linkage Map of the Zebra Finch Taeniopygia guttata Provides New Insights Into Avian Genome Evolution. Genetics, 179(1): 651–667.

Stapley, J., Feulner, P. G. D., Johnston, S. E., Santure, A. W., and Smadja, C. M. 2017. Variation in recombination frequency and distribution across eukaryotes: patterns and processes. Phil. Trans. R. Soc. B, 372(1736): 20160455.

Van der Auwera, G. A., Carneiro, M. O., Hartl, C., Poplin, R., Del Angel, G., Levy-Moonshine, A., Jordan, T., Shakir, K., Roazen, D., Thibault, J., Banks, E., Garimella, K. V., Altshuler, D., Gabriel, S., and DePristo, M. A. 2013. From FastQ data to high confidence variant calls: the Genome Analysis Toolkit best practices pipeline. Current Protocols in Bioinformatics / Editoral Board, Andreas D. Baxevanis … [et Al.], 43: 11.10.1–33.

van Oers, K., Santure, A. W., De Cauwer, I., van Bers, N. E., Crooijmans, R. P., Sheldon, B. C., Visser, M. E., Slate, J., and Groenen, M. A. 2014. Replicated high-density genetic maps of two great tit populations reveal fine-scale genomic departures from sex-equal recombination rates. Heredity, 112(3): 307–316.

Wallberg, A., Glémin, S., and Webster, M. T. 2015. Extreme Recombination Frequencies Shape Genome Variation and Evolution in the Honeybee, Apis mellifera. PLOS Genetics, 11(4): e1005189.

Warren, W. C., Clayton, D. F., Ellegren, H., Arnold, A. P., Hillier, L. W., Künstner, A., Searle, S., White, S., Vilella, A. J., Fairley, S., Heger, A., Kong, L., Ponting, C. P., Jarvis, E. D., Mello, C. V., Minx, P., Lovell, P., Velho, T. A. F., Ferris, M., Balakrishnan, C. N., Sinha, S., Blatti, C., London, S. E., Li, Y., Lin, Y.-C., George, J., Sweedler, J., Southey, B., Gunaratne, P., Watson, M., Nam, K., Backström, N., Smeds, L., Nabholz, B., Itoh, Y., Whitney, O., Pfenning, A. R., Howard, J., Völker, M., Skinner, B. M., Griffin, D. K., Ye, L., McLaren, W. M., Flicek, P., Quesada, V., Velasco, G., Lopez-Otin, C., Puente, X. S., Olender, T., Lancet, D., Smit, A. F. A., Hubley, R., Konkel, M. K., Walker, J. A., Batzer, M. A., Gu, W., Pollock, D. D., Chen, L., Cheng, Z., Eichler, E. E., Stapley, J., Slate, J., Ekblom, R., Birkhead, T., Burke, T., Burt, D., Scharff, C., Adam, I., Richard, H., Sultan, M., Soldatov, A., Lehrach, H., Edwards, S. V., Yang, S.-P., Li, X., Graves, T., Fulton, L., Nelson, J., Chinwalla, A., Hou, S., Mardis, E. R., and Wilson, R. K. 2010. The genome of a songbird. Nature, 464(7289): 757–762.

Weber, C. C., Boussau, B., Romiguier, J., Jarvis, E. D., and Ellegren, H. 2014. Evidence for GC-biased gene conversion as a driver of between-lineage differences in avian base composition. Genome Biology, 15(12): 549.

Williams, A. L., Genovese, G., Dyer, T., Altemose, N., Truax, K., Jun, G., Patterson, N., Myers, S. R., Curran, J. E., Duggirala, R., Blangero, J., Reich, D., and Przeworski, M. 2015. Non-crossover gene conversions show strong GC bias and unexpected clustering in humans. eLife, 4: e04637.

Yang, Z. 2007. PAML 4: Phylogenetic Analysis by Maximum Likelihood. Molecular Biology and Evolution, 24(8): 1586–1591.

Zeng, K., Jackson, B. C., and Barton, H. J. 2019. Methods for Estimating Demography and Detecting Between-Locus Differences in the Effective Population Size and Mutation Rate. Molecular Biology and Evolution, 36(2): 423–433.

Zhang, G., Li, C., Li, Q., Li, B., Larkin, D. M., Lee, C., Storz, J. F., Antunes, A., Greenwold, M. J., Meredith, R. W., Ödeen, A., Cui, J., Zhou, Q., Xu, L., Pan, H., Wang, Z., Jin, L., Zhang, P., Hu, H., Yang, W., Hu, J., Xiao, J., Yang, Z., Liu, Y., Xie, Q., Yu, H., Lian, J., Wen, P., Zhang, F., Li, H., Zeng, Y., Xiong, Z., Liu, S., Zhou, L., Huang, Z., An, N., Wang, J., Zheng, Q., Xiong, Y., Wang, G., Wang, B., Wang, J., Fan, Y., da Fonseca, R. R., Alfaro-Nuñez, A., Schubert, M., Orlando, L., Mourier, T., Howard, J. T., Ganapathy, G., Pfenning, A., Whitney, O., Rivas, M. V., Hara, E., Smith, J., Farré, M., Narayan, J., Slavov, G., Romanov, M. N., Borges, R., Machado, J. P., Khan, I., Springer, M. S., Gatesy, J., Hoffmann, F. G., Opazo, J. C., Håstad, O., Sawyer, R. H., Kim, H., Kim, K.-W., Kim, H. J., Cho, S., Li, N., Huang, Y., Bruford, M. W., Zhan, X., Dixon, A., Bertelsen, M. F., Derryberry, E., Warren, W., Wilson, R. K., Li, S., Ray, D. A., Green, R. E., O’Brien, S. J., Griffin, D., Johnson, W. E., Haussler, D., Ryder, O. A., Willerslev, E., Graves, G. R., Alström, P., Fjeldså, J., Mindell, D. P., Edwards, S. V., Braun, E. L., Rahbek, C., Burt, D. W., Houde, P., Zhang, Y., Yang, H., Wang, J., Avian Genome Consortium, Jarvis, E. D., Gilbert, M. T. P., and Wang, J. 2014. Comparative genomics reveals insights into avian genome evolution and adaptation. Science (New York, N.Y.), 346(6215): 1311–1320.

