## Supplementary figures and tables for "The effective population size modulates the strength of GC biased gene conversion in two passerines"

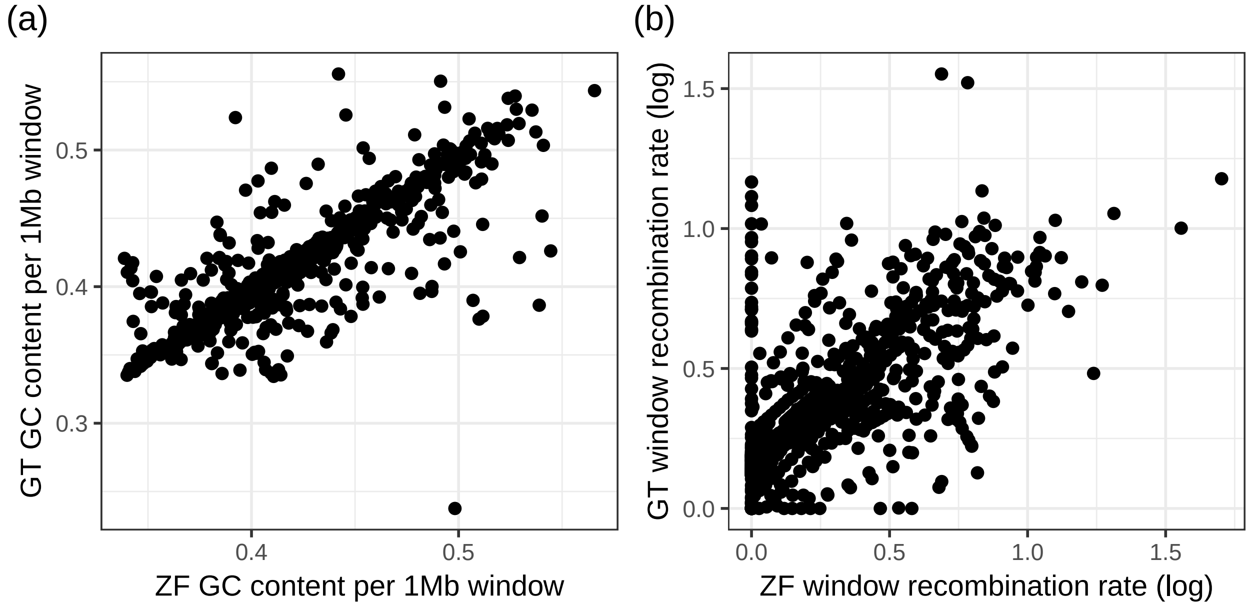


Figure S1: Relationships between zebra finch GC content and great tit GC content (a) and zebra finch recombination rate and great tit recombination rate (b) across the 1Mb window dataset.

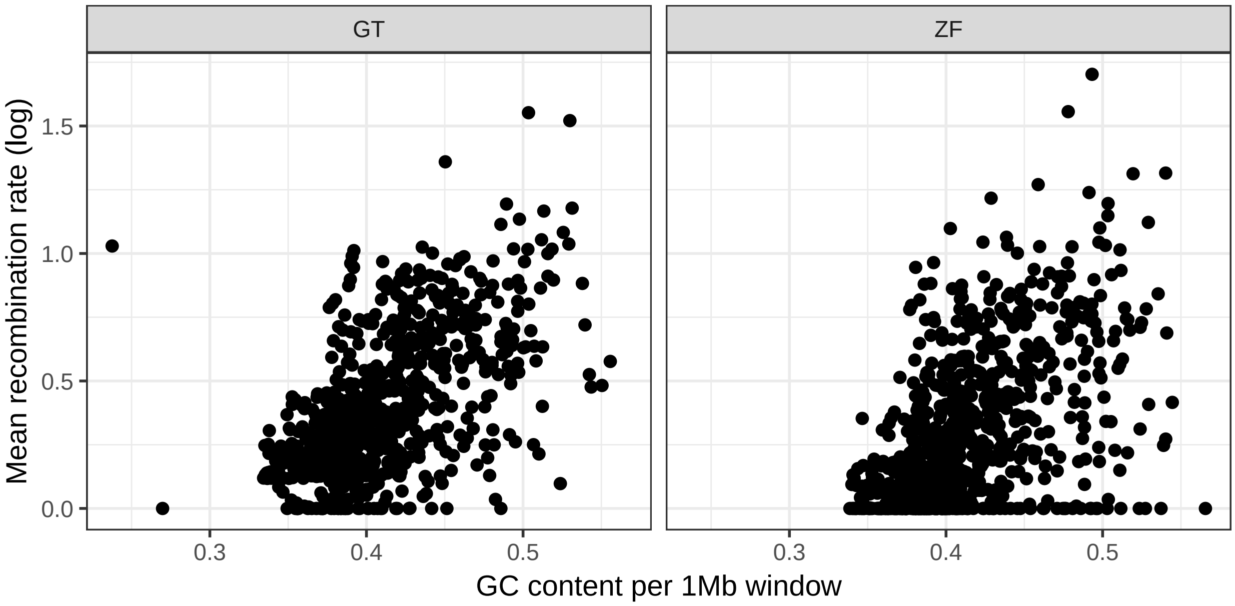


Figure S2: Relationships between GC content and recombination rate across the 1Mb window dataset in both the great tit (GT) and the zebra finch (ZF).

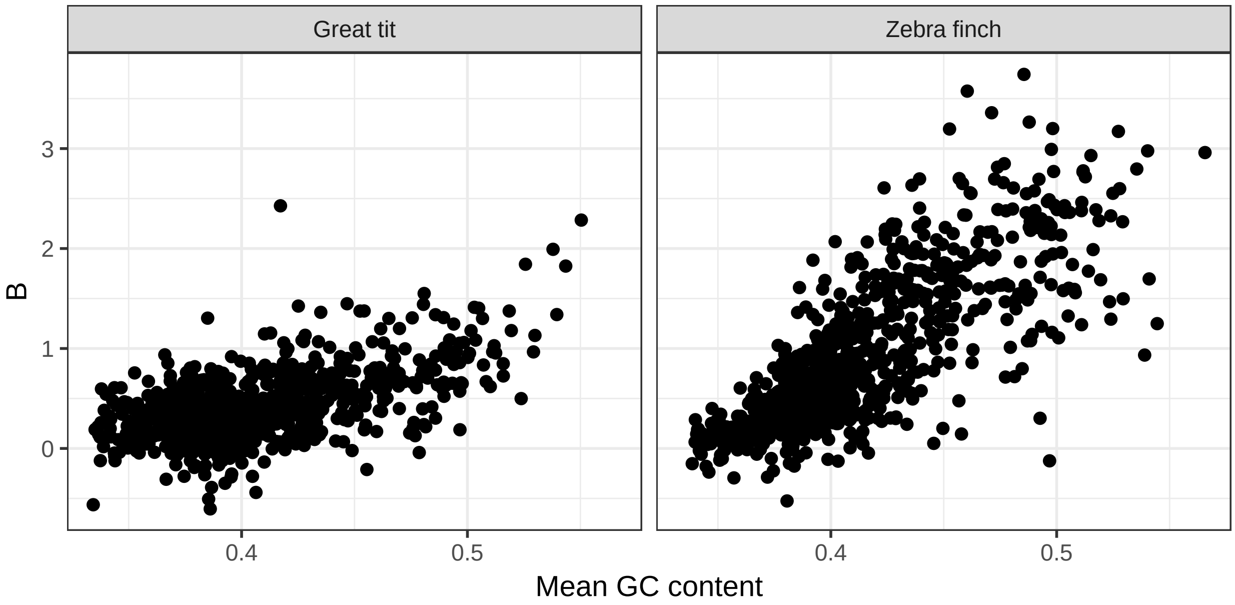


Figure S3: The relationship between mean window GC content and the strength of gene conversion (B) in the great tit and zebra finch.

Table S1: Results of multiple regression analysis of the strength of gene conversion ($B$) against GC content and $\pi$, and against recombination rate and $\pi$, separately, using a dataset with CpG sites removed for both species. Importance is the relative importance (as a proportion of the total variance explained) as estimated using the pmvd method implemented in the `relaimpo` package (Groemping, 2006).

| model | species | variable | estimate | importance | p value | $R^2$ |
|---|---|---|---|---|---|---|
| $B \sim$ log10(crossover rate + 1) + $\pi$ | great tit | crossover rate | 0.816 | 0.91 | $< 2 \times 10^{-16}$ | 0.336 |
| | | $\pi$ | 112 | 0.09 | $3.81 \times 10^{-9}$ | |
| | zebra finch | crossover rate | 1.78 | 0.85 | $< 2 \times 10^{-16}$ | 0.458 |
| | | $\pi$ | 102 | 0.15 | $< 2 \times 10^{-16}$ | |
| $B \sim$ GC content + $\pi$ | great tit | GC content | 6.01 | 0.80 | $< 2 \times 10^{-16}$ | 0.414 |
| | | $\pi$ | 201 | 0.20 | $< 2 \times 10^{-16}$ | |
| | zebra finch | GC content | 16.4 | 0.95 | $< 2 \times 10^{-16}$ | 0.623 |
| | | $\pi$ | 73.5 | 0.05 | $5.32 \times 10^{-15}$ | |

Table S2: GC content in coding and non-coding regions in the great tit and zebra finch genomes.

| region | species | GC |
|---|---|---|
| non-coding | great tit | 0.41 |
| coding | great tit | 0.52 |
| non-coding | zebra finch | 0.41 |
| coding | zebra finch | 0.48 |

Table S3: Maximum likelihood parameter estimates from fitting a 2 epoch demographic model to weak to weak and strong to strong (GC conservative) SNPs using VarNe for both the great tit and the zebra finch. The parameters are as follows, the population scaled mutation rate: $\theta = 4N_e\mu$, the relative size of the population size change: $g$, the timing of the population size change in units of $2N_e$: $\tau$, and the polarisation error rate: $\epsilon$. $\tau$ is also reported re-scaled to years with $years = \frac{\tau\theta T_{gen}}{2\mu_{wwss}}$, where $T_{gen}$ is the generation time and $\mu_{wwss}$ is the per site per generation mutation rate for GC conservative SNPs, taken to be $\mu/3$, using the estimates in table S5. Values in bracket represent the 95% confidence intervals resulting from 100 rounds of resampling with replacement.

| species | $\theta$ | $g$ | $\tau$ | $\epsilon$ | years |
|---|---|---|---|---|---|
| zebra finch | 0.000607 | 12.3 | 1.25 | 0.0186 | 494837 |
| | (0.000593, 0.000620) | (11.9, 12.8) | (1.21, 1.30) | (0.0181, 0.0190) | (467954, 525652) |
| great tit | 0.000448 | 1.89 | 0.208 | 0.0562 | 139776 |
| | (0.000439, 0.000456) | (1.86, 1.92) | (0.199, 0.219) | (0.0547, 0.0573) | (131042, 149796) |

Table S4: Mean maximum likelihood parameter estimates from fitting a 2 epoch demographic model to weak to weak and strong to strong (GC conservative) SNPs using VarNe to each 1Mb window in our dataset, for both the great tit and the zebra finch. The parameters are as follows, the population scaled mutation rate: $\theta = 4N_e\mu$, the relative size of the population size change: $g$, the timing of the population size change in units of $2N_e$: $\tau$, and the polarisation error rate: $\epsilon$.

| species | $\theta$ | $g$ | $\tau$ | $\epsilon$ |
|---|---|---|---|---|
| zebra finch | 0.00108 | 15.8 | 1.51 | 0.0219 |
| great tit | 0.00108 | 2.70 | 0.373 | 0.0569 |

Table S5: $N_e$ estimates for each species, calculated using published estimates of nucleotide diversity at fourfold degenerate sites ($\pi_4$) and the per year per site mutation rate ($\mu$). Calculated with $N_e = \pi_4/4\mu$. Generation times and $\mu$ estimates were used to re-scale times to years in the demographic analyses.

| species | $\pi_4$ | $\mu$ | $N_e$ | generation time |
|---|---|---|---|---|
| zebra finch | 0.0099 | $2.3 \times 10^{-9}$ | $1,076,087$ | 1 year |
| | (Corcoran *et al.*, 2017) | Smeds *et al.* (2016) | | Shultz *et al.* (2016) |
| great tit | 0.0035 | $2 \times 10^{-9}$ | $437,500$ | 2 year |
| | (Corcoran *et al.*, 2017) | Laine *et al.* (2016) | | Laine *et al.* (2016) |

### References


Corcoran, P., Gossmann, T. I., Barton, H. J., Great Tit HapMap Consortium, Slate, J., and Zeng, K. 2017. Determinants of the Efficacy of Natural Selection on Coding and Noncoding Variability in Two Passerine Species. *Genome Biol Evol*, 9(11): 2987–3007.

Groemping, U. 2006. Relative Importance for Linear Regression in R: The Package relaimpo. *Journal of Statistical Software*, 17(1): 1–27. Number: 1.

Laine, V. N., Gossmann, T. I., Schachtschneider, K. M., Garroway, C. J., Madsen, O., Verhoeven, K. J. F., de Jager, V., Megens, H.-J., Warren, W. C., Minx, P., Crooijmans, R. P. M. A., Corcoran, P., Great Tit HapMap Consortium, Sheldon, B. C., Slate, J., Zeng, K., van Oers, K., Visser, M. E., and Groenen, M. A. M. 2016. Evolutionary signals of selection on cognition from the great tit genome and methylome. *Nat Commun*, 7: 10474.

Shultz, A. J., Baker, A. J., Hill, G. E., Nolan, P. M., and Edwards, S. V. 2016. SNPs across time and space: population genomic signatures of founder events and epizootics in the House Finch (Haemorhous mexicanus). *Ecology and Evolution*, 6(20): 7475–7489. _eprint: https://onlinelibrary.wiley.com/doi/pdf/10.1002/ece3.2444.

Smeds, L., Qvarnström, A., and Ellegren, H. 2016. Direct estimate of the rate of germline mutation in a bird. *Genome Research*, 26(9): 1211–1218.
